# Extracellular Vesicle-Associated Neutrophil Elastase Activates Hepatic Stellate Cells and Promotes Liver Fibrogenesis via ERK1/2 Pathway

**DOI:** 10.1101/2024.08.20.608832

**Authors:** Regina Oshins, Zachary Greenberg, Yun-Ling Tai, Derrick Zhao, Xuan Wang, Borna Mehrad, Mei He, Ishan Patel, Laith Khartabil, Huiping Zhou, Mark Brantly, Nazli Khodayari

## Abstract

Liver fibrosis associated with increased mortality is caused by activation of hepatic stellate cells and excessive production and accumulation of extracellular matrix in response to fibrotic insults. It has been shown that in addition to liver inflammation, systemic inflammation also contributes to liver fibrogenesis. A deeper understanding of mechanisms that control liver fibrotic response to intra-and extra-hepatic inflammation is essential to develop novel clinical strategies against this disease. Extracellular vesicles (EV) have been recognized as immune mediators that facilitate activation of hepatic stellate cells. In inflammatory diseases, activated neutrophils release neutrophil elastase (NE) bound to EV, which has been identified as a significant contributor to inflammation by promoting immune cell activation. Here, we aimed to explore the role of inflammation derived plasma EV-associated NE in liver fibrogenesis and its potential mechanisms. We show EV-associated NE induces activation, proliferation and migration of hepatic stellate cells by promoting activation of the ERK1/2 signaling pathway. This effect did not occur through EV without surface NE, and Sivelestat, a NE inhibitor, inhibited activation of the ERK1/2 signaling pathway mediated by EV-associated NE. Moreover, we found plasma EV-associated NE increases deposition of collagen1 and α-smooth muscle actin in the liver of a mouse model of liver fibrosis (Mdr2^-/-^). Notably, this effect does not occur in control mice without preexisting liver disease. These data suggest that EV-associated NE is a pro-fibrogenic factor for hepatic stellate cell activation via the ERK1/2 signaling pathway in pre-existing liver injuries. Inhibition of the plasma EV-associated NE in inflammatory conditions may be a therapeutic target for liver fibrosis in patients with inflammatory diseases.

## Introduction

Liver fibrosis is a severe global health issue in individuals with chronic liver disease with potential to progress to cirrhosis and hepatocellular carcinoma (1). Excessive accumulation of extracellular matrix (ECM) is a hallmark of liver fibrosis. Activated hepatic stellate cells (HSC) are the primary source of extracellular matrix proteins in all types of fibrotic liver diseases (2). Recent studies show HSC are responsive to extracellular signals derived from immune cells and exert an effect on regulation of fibrotic response of the liver during inflammation (3). Systemic inflammation is implicated in activation of HSC in chronic liver diseases (4). Given this, targeting systemic inflammation and its role in liver fibrogenesis could offer therapeutic strategies for managing fibrosis and related conditions.

Neutrophils are the most abundant leukocytes, playing an important role in the innate immune system (5, 6). Neutrophils are persistently recruited to sites of inflammation, releasing cytokines and proteases to activate immune cells and regulate inflammation (7). Neutrophil elastase (NE) plays an important pathologic role in a variety of chronic inflammatory diseases, such as lung and liver inflammation (8). Experimental studies demonstrate NE plays a significant role in inflammatory responses by activating surface receptors and signaling pathways, leading to cellular activation at inflammation sites (9, 10). The imbalance between NE and protease inhibitors is implicated in the pathogenesis of several inflammatory diseases, such as lung and liver ECM remodeling (11, 12).

Extracellular vesicles (EV), ranging from 50-150 nm in diameter, are membrane-bound vesicles released by most cell types under physiologic conditions and in response to stimuli (13). Recent work has shown EV released from activated neutrophils contain NE with intact enzymatic activity which is resistant to protease inhibitors such as α1-antitrypsin (14, 15). EV positive for NE (NE^+^EV) exist in plasma from patients with lung disease or systemic inflammation and these EV are capable of transferring inflammation to mice in an NE-dependent manner (15). However, the relationship between liver fibrosis and systemic inflammation in the context of plasma circulating NE^+^EV remains to be explored.

This study was designed to investigate the link between NE^+^EV and liver fibrogenesis with a focus on regulating HSC activation. Our findings reveal EV-associated NE triggers HSC activation through the ERK1/2 signaling pathway, both *in vitro* and *in vivo*. Additionally, NE^+^EV were shown to exacerbate liver fibrosis in the Mdr2^-/-^ mouse model of spontaneous liver fibrosis. This research highlights a novel mechanism by which systemic inflammation drives progression of liver fibrosis in patients with liver disease. These results suggest targeting NE^+^EV could be a potential strategy to mitigate liver fibrosis in patients with systemic inflammation.

## Methods

### Mice and treatments

Six-to-eight-week-old male and female C57BL/6 mice were purchased from The Jackson Laboratory. All animals were maintained under 12 h light/dark conditions at 22– 24°C with unrestricted access to food and water. All animal protocols were performed according to guidelines for care and use of laboratory animals and approved by the animal ethics committee of their respective universities. To induce lung inflammation, mice were randomly divided into groups: control (intranasal administration of saline at 50μL/mouse), and experimental (intranasal administration of LPS at 40µg/mouse). Five mice were included per group. After 24 hours, samples of bronchoalveolar lavage fluid (BALF), lung homogenate and plasma were collected for further analysis.

To induce liver fibrosis, eight-week-old male and female Mdr2^-/-^ mice with FVB background were treated via tail vein injection twice weekly for four weeks with 10^9^ EV collected from either quiescent or n-fMLP activated mouse neutrophils. Mdr2^+/-^ mice were used as a control. At the end of treatment, mice were sacrificed, and blood and liver tissues were harvested for further analysis.

### Murine bronchoalveolar lavage fluid

A midline incision was performed from the upper abdomen to the mid-cervical region, exposing the lungs and trachea of mouse. An 18-gauge venous catheter was inserted into the cervical trachea and sealed around the cannula with gently tied suture.

Bronchoalveolar lavage (BAL) was then performed by gently instilling 2mL of PBS over 2 periods of 1 minute each, followed by slow retrieval of infused fluid into a syringe.

### Preparation of mouse neutrophils from the bone marrow

Bone marrow was collected from mouse tibias and femurs by centrifugation. Cells were pelleted into 50μL of RPMI containing 2% FBS and 1% penicillin/streptomycin. Red cells were lysed for 1 minute at room temperature in ACK buffer. Cells were then centrifuged for 5 minutes at 400xg and 4°C. Cells were again suspended in RPMI, washed through a 100μm cell strainer and centrifuged for 5 minutes at 400xg and 4°C. Bone marrow cells then were suspended in 3mL of cold, sterile PBS. A gradient is formed by placing 3mL of Histopaque 1119 (Sigma, St Louis, MO) in a 15mL conical tube, overlaying 3mL of Ficol (Cytiva, Marlborough, MA), and then overlaying the cells in PBS. The gradient was centrifuged for 30 minutes at 870xg at room temperature without brakes. Neutrophils were collected from the Histopaque/Ficol interface and washed twice with PBS.

### Neutrophil stimulation

Neutrophils were suspended in RPMI containing 5% exosome-free FBS and 1% penicillin/streptomycin. For activation of the neutrophils, the culture media also contains 2μM n-fMLP (MilliporeSigma, Burlington, MA). Cells were incubated 30 minutes at 37°C and 5% CO_2_.

EV isolation and characterization

### Differential ultracentrifugation

EV were isolated from cell culture media and bronchioalveolar lavage (BAL) using differential centrifugation as previously described (16). Briefly, samples were ultracentrifuged in a Beckman Coulter Optima XE90 Ultracentrifuge with an SW-40Ti swinging rotor at 110,000xg for 2 hours at 4°C. EV pellets were resuspended in PBS, filtered through a 0.22μm filter (Millipore, Billerica, MA) and centrifuged at 110,000xg for 70 minutes at 4°C. EV were resuspended in 200μL of PBS and conserved until further experiments. For NE removal experiments, activated neutrophil-derived EVs were pre-incubated with 25μM protamine sulfate (CalBiochem) for 30 minutes at room temperature prior to filtration.

### NanoPom NE^+^EV and NE^-^EV purification

NE^+^EV positive (CD9/CD63/CD81^+^) and NE^-^EV (CD9/CD63/CD81^+^) were isolated using NanoPom as previously described (17). In brief, 10mL of cell culture media was sequentially centrifuged at 2000xg, then 10000xg, for 15 minutes each, to obtain pre-cleared supernatant. 40μL of NanoPom beads utilizing anti-Neutrophil Elastase (ThermoScientific, Waltham, MA) were injected into the media to capture NE^+^EV. After 24 hours, captured NE^+^EV were magnetically separated, washed 3 times with ice-cold PBS, then released into 200μL. Next, NanoPom beads utilizing anti-CD9/CD63/CD81 to represent NE^-^EV were injected into the same media, followed by the same procedure to capture, wash and release EV into 200μL for downstream assays.

### Nanoparticle Tracking Analysis

Nanoparticle tracking analysis was conducted using ZetaView (QUATT, Particle Metrix Inc, USA). ZetaView measures the nanoparticle’s Brownian motion using an incident laser to determine its corresponding size. The nanoparticle motion is then tracked by the detector and recorded over time. The incident laser wavelength was 488 nm^-1^ with sensitivity at 75 and shutter time at 163, over 90 seconds at the highest video resolution for all 11 positions. The size and number of purified EV were then quantified by injecting 100μL of isolated EV into 900μL of 1x PBS into the ZetaView.

### Cell culture and treatments

LX2 cells (a human HSC line) were purchased from MilliporeSigma. Cells were cultured in DMEM containing 2% FBS, 1% penicillin/streptomycin and 1X glutamine (Invitrogen, Waltham, MA) at 37°C and 5% CO_2_. When LX2 cells reached 60% confluence, they were passaged into low glucose DMEM containing 1X serum replacement 1 (Sigma), 100μM vitamin A, 50ng/mL insulin, 0.5mM glutamine and 1% penicillin/streptomycin and treated with EV (10^9) for indicated times.

### Migration assay

LX2 cells were plated at 500,000 cells/well in 6 well plates and grown for 16h. The medium was aspirated, and the cell-coated surface was scraped with a 200μL pipette tip in a single stripe. The scrape-wounded surface was washed twice with PBS and the cultures were treated with and without NE^+^EV and NE^-^EV and allowed to heal for 30h at 37 °C. Migration of cells into wounded areas was evaluated with an inverted microscope and photographed. The average extent of wound closure was evaluated by multiple measurements of the width of the wound space for each case.

### Gel contraction assay

Matrigel (Corning, USA) was added to 24-well plates and incubated at room temperature for 1 hour to allow gelation. LX2 cells were seeded on the Matrigel at equal density in each well and incubated at 37°C to adhere for two hours. LX2 cells were treated with NE^+^EV or NE^-^EV and the gels detached from the walls of the cell culture plate using a pipet tip. Over a 48-hour period, images were taken to monitor the changes of the gel. Measurement of the gel area was quantified using Image J.

### Transmission electron microscopy

Purified EV were fixed with 2% paraformaldehyde. 20μL of the EV suspension was loaded onto a formvar coated grid, negatively stained with 2% aqueous uranyl acetate, and examined under a Hitachi 7600 transmission electron microscope (Hitachi High-Technologies, Schaumburg, IL) equipped with a Macrofire monochrome progressive scan CCD camera (Optronics, Goleta, CA) and AMTV image capture software (Advanced Microscopy Techniques, Danvers, MA) (16).

### DIR fluorescent labeling of EV

EV were stained with DiR (Invitrogen, Waltham, MA) in the dark at a concentration of 5μg/mL for 30 minutes at room temperature. PBS mixed with the same concentration of DiR serves as a control. After staining, samples were washed with PBS and pelleted by ultracentrifugation.

### Western blot analysis

LX2 cells were seeded at 5×10^5^/well in 6-well plates with or without NE^-^EV or NE^+^EV for indicated times. Protein levels were determined in the whole cell or EV lysate homogenates in RIPA buffer using the bicinchoninic acid method (Pierce Biotechnology, Rockford, IL). Rabbit polyclonal antibodies were used to detect total and phospho-ERK1/2, total and phospho-MEK, Calnexin, Actin (Cell Signaling, Danvers, MA), CD81, NE, TSG101 and GAPDH (Proteintech, Rosemont, IL) (16). Proteins were detected using Super Signal West Dura Extended Duration Substrate Kit (ThermoScientific, Waltham, MA).

### Immunofluorescence and immunohistochemistry

For immunohistochemical staining, paraffin-embedded liver tissue sections were sliced into 5μm sections, de-paraffinized with xylene and rehydrated with an ethanol gradient. Antigen retrieval was achieved by a 20-minute incubation in 95°C sodium citrate buffer (pH 6.0) and 20 minutes of cooling at room temperature. Endogenous peroxidases were quenched by incubation in 3% hydrogen peroxide for 20 minutes. Sections were washed with PBS and primary antibodies applied for 60 minutes at room temperature in a humidified chamber. After washing in PBS, slides were incubated in secondary antibody for 1 hour at room temperature. After washing in PBS, slides were incubated with Vectastain ABC (Vector Laboratories) for 30 minutes. After washing in PBS, color development was achieved with diaminobenzidine tetrahydrochloride (DAB) (Vector Laboratories) for two to five minutes.

For immunofluorescent staining, paraffin-embedded liver tissue sections were sliced into 5μm sections, de-paraffinized with xylene and rehydrated with an ethanol gradient. Epitope retrieval was performed for 30 minutes in a steam chamber using Epitope Retrieval Solution (IHC World, Ellicott City, MD) after which the slides were allowed to cool in the solution for 15 minutes at room temperature before being washed. Slides were blocked in 2.5% normal goat serum for 20 minutes at room temperature and incubated with phospho-ERK (Cell Signaling, Danvers, MA) and Vimentin (ThermoScientific, Waltham, MA) primary antibodies overnight at 4°C. After washing, slides were incubated for 45 minutes at room temperature with secondary antibodies. pERK was conjugated with goat anti-rabbit 594 and vimentin with goat anti-rat 488 (ThermoScientific, Waltham, MA). After washing, slides were stained for 10 minutes at room temperature with 2 μg/mL Hoechst. Images were acquired and signal intensity quantified using a Keyence BZ-X700 microscope.

### Enzyme-linked immunosorbent assay (ELISA)

NE was measured by ELISA following manufacturer’s instructions (Abcam, Waltham, MA). Plasma EV samples were first lysed in an equal volume of RIPA buffer and then diluted in sample buffer to a final dilution of 1:10. BAL EV were lysed in an equal volume of RIPA buffer and then diluted in sample buffer to a final concentration of 1:5. Plasma supernatant was directly diluted at 1:25 and BAL supernatant at 1:5 in sample buffer. Samples were incubated with antibody cocktail for 1 hour at room temperature in the assay plate and developed with TMB substrate for 10 minutes before the optical density was measured using a SpectraMax spectrometer at 450 nm and the sample concentrations calculated based on the standard curve.

### Flow Cytometry

Lung nonparenchymal cells isolated from WT mice with and without LPS treatment (*n=*5/group) were stained as previously described (18) and analyzed using a CytoFlex (Beckman Coulter, Brea, CA). Dead cells were excluded using Live/Dead Near IR (Invitrogen, Carlsbad, CA). Cells were stained with CD45-PerCP, CD11b-BV750, Ly6C-PE/Cy7 and Ly6G-BV605 antibodies from BioLegend (San Jose, CA). Data were analyzed using FlowJo (BD Biosciences, San Jose, CA) software.

### RNA Extraction and Quantitative Real-Time PCR

Total RNA was extracted using a Qiagen RNeasy Plus Mini Kit and reverse transcribed into cDNA using SuperScript VILO Master Mix. Gene expression levels were analyzed by qPCR using an Applied Biosystems 7500 fast real-time PCR system and Taqman Fast Advanced Master Mix. Relative expression levels of target mRNAs were determined using TaqMan probes normalized with 18S as an endogenous control and analyzed by the 2^-ΔΔCT^ method as previously described (16).

## Statistical analysis

All results and data presented in this study are expressed as means ± SD. Statistical analyses were performed using the Prism9 software program (GraphPad Software) by Student’s t test or Mann–Whitney U test. One-way ANOVA test followed by Newman-Keuls test was performed for the multiple-group comparisons. Non-parametric Kruskal-Wallis test was used for data that cannot pass the normality test. Values of p<0.05 were considered statistically significant.

## Results

### Characterization of EV fraction

EV were isolated from both activated and quiescent bone marrow-derived neutrophils and from BALF using a combination of filtration, ultracentrifugation and NanoPom (Figure 1A, BioRender.com). Characterization of EV was performed by analyzing particle size and concentration using Nanoparticle Tracking Analysis (NTA), revealing all EV were within the expected range of 100–150 nm in diameter (Figure 1B). The yield of EV from cultured media of activated neutrophils was comparable to that from quiescent neutrophils (Figure 1C), and the concentration of EV correlated with the number of cultured neutrophils (Figure 1D). Transmission Electron Microscopy (TEM) with negative staining showed cup-shaped, round particles, confirming typical morphology of isolated EV from mouse BAL (Figure 1E). Next, we performed western blot analysis to validate the presence of EV markers in mice BAL derived EV. Our results demonstrated abundant expression of CD81, TSG101 and actin in all EV fractions. Calnexin was used as a negative control for EV and was absent, confirming the specificity of the EV isolation (Figure 1F).

**Figure 1.**
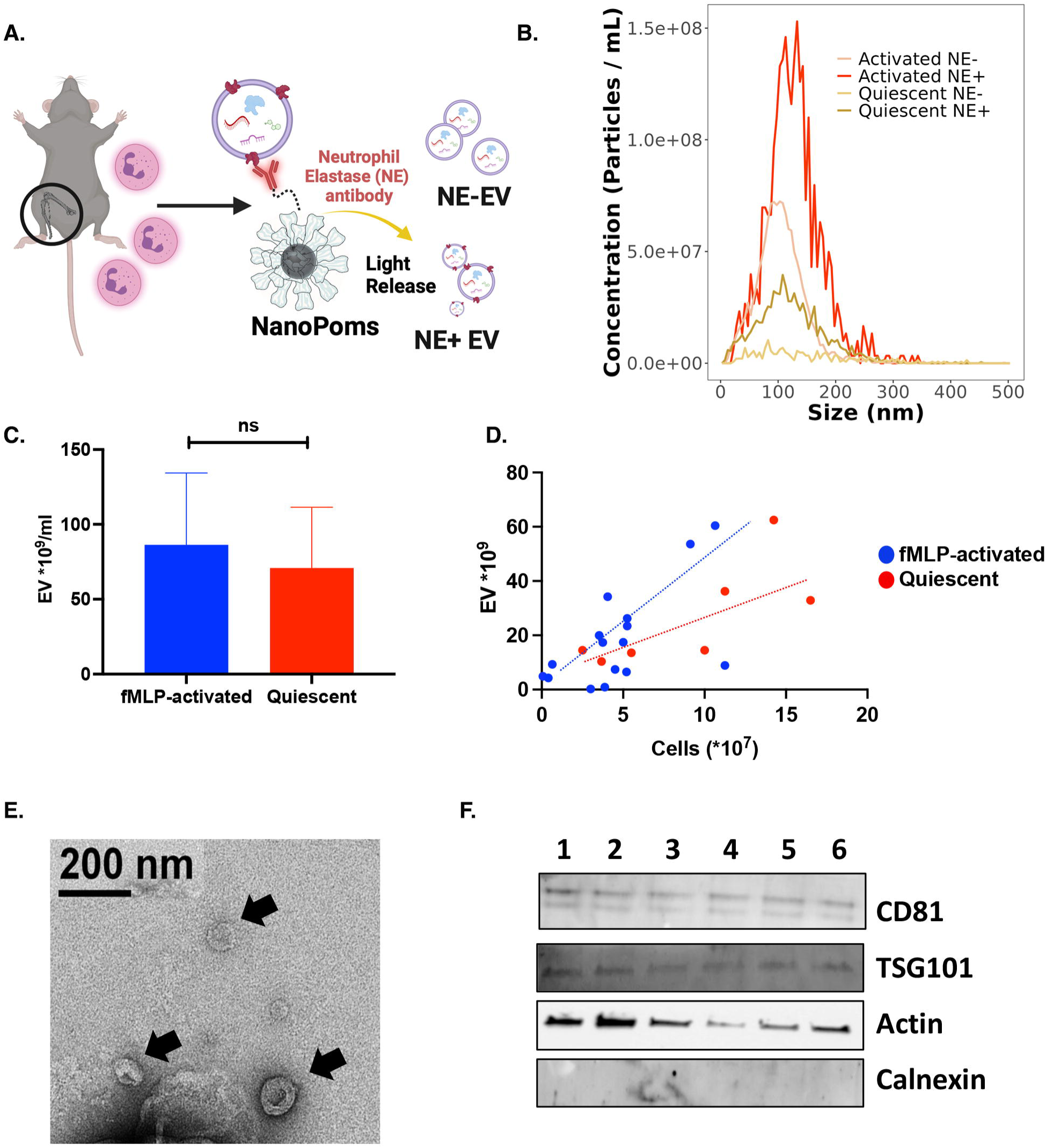
Characterization of isolated extracellular vesicles (EV). **(A)** Schematic presentation of EV isolation derived from bone marrow neutrophils. Neutrophils were isolated from the tibia and femur of wildtype mice, cultured and activated. EV were isolated from activated or quiescent neutrophils using a combination of filtration and ultracentrifugation, or NanoPom technology (BioRender.com). **(B)** Nanoparticle Tracking Analysis (NTA) was used to characterize EV particle size and concentration. The results from NTA showed all EV are within the normal range for EV (100–150 nm in diameter). NTA also revealed that activated neutrophils release a higher number of NE+EV compared to quiescent neutrophils. **(C)** While the EV recovery from activated and quiescent neutrophil culture media showed no significance differences, (**D)** the concentration of isolated EV from the culture media of both activated and quiescent neutrophils were correlated with the number of cultured neutrophils. **(E)** Negative staining of mouse BALF-derived EV using Transmission Electron Microscopy (TEM) demonstrated cup-shaped, round particles for EV fraction. **(F)** Western blot analysis was used to investigate the presence of EV markers in 6 different mice BALF-derived EV fractions. The results confirmed presence of CD81, TSG101 and actin in all EV fractions. Calnexin was used as EV negative control marker.

Lung Inflammation induced by LPS Increases BALF and Plasma Circulating NE^+^EV *In-vivo* Given higher numbers of NE^+^EV in mouse BALF during lung inflammation (15, 19), we examined the presence of NE^+^EV in the circulating blood of the mice 24 h after LPS instillation (40µg/mouse) induced lung inflammation (n=5/group). To assess lung infiltration of neutrophils in response to LPS, we characterized immune cell populations in lungs of mice. Neutrophils were identified by expression of CD45, CD11b and Ly6G and lack of Ly6C (Figure 2A). Our results showed lung inflammatory response to LPS, indicated by increased neutrophil population in lungs of LPS treated mice as compared to controls (Figure 2B). Next, EV were isolated from BALF and serum. Levels of NE were measured from the EV fraction and EV-depleted BALF and serum of mice using ELISA. We observed that LPS-driven lung injury increases levels of both EV-associated NE and free NE in the BALF of LPS treated as compared to control mice (Figure 2C). However, our results showed LPS-driven lung injury specifically increased levels of EV-associated NE in plasma of LPS-treated mice. There were no significant differences in levels of free NE in EV-depleted serum between the two groups (Figure 2D).

**Figure 2.**
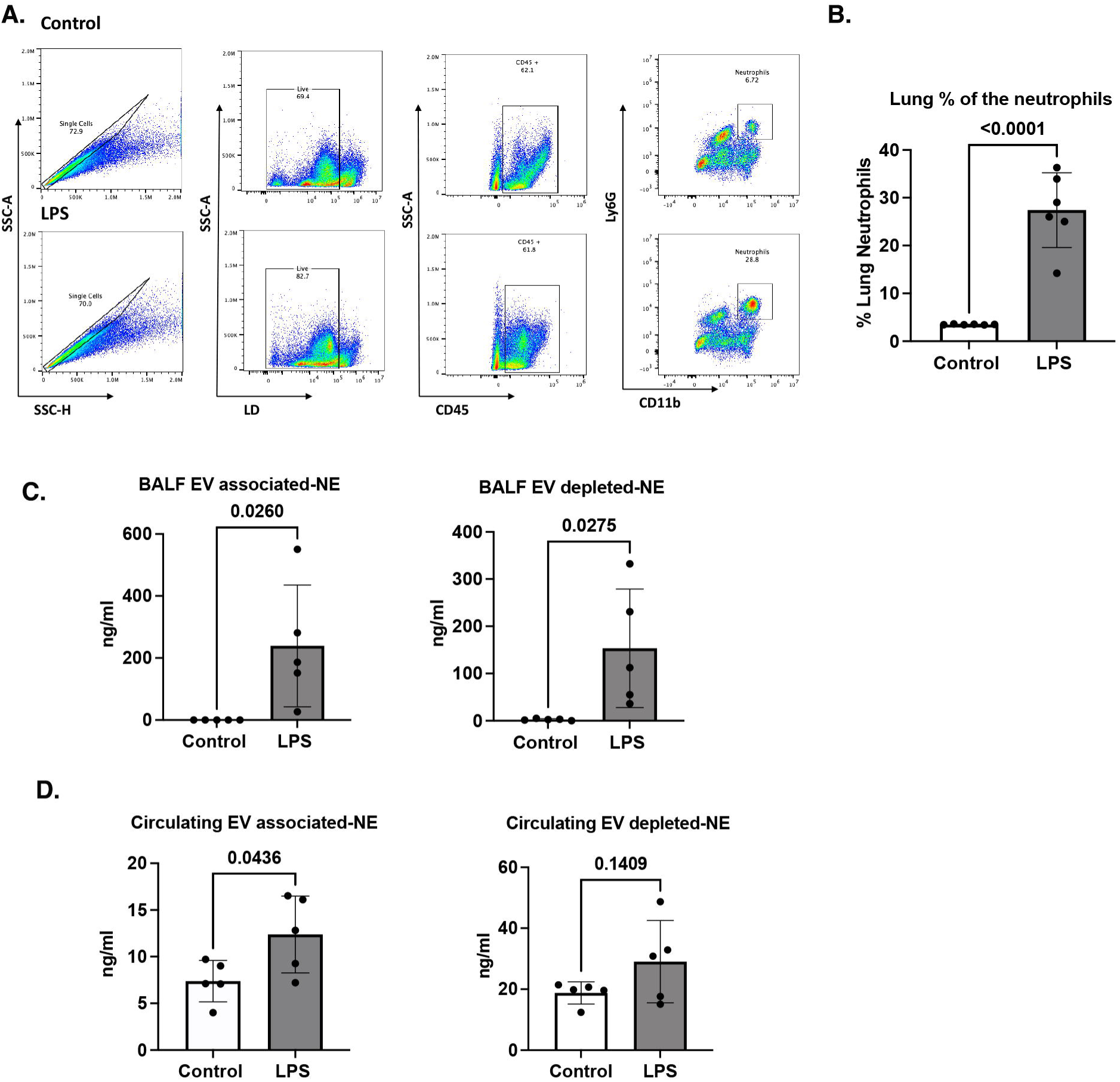
Lung Inflammation results in an increase in NE^+^EV both in BALF (bronchioalveolar lavage fluid) and plasma. **(A)** Flow cytometry was used to assess lung infiltration of neutrophils in response to LPS induced inflammation in mice. Neutrophils were identified by expression of CD45, CD11b and Ly6G and lack of Ly6C markers. **(B)** Flow cytometry results showed lung LPS administration results in increased neutrophil population in the lungs of LPS treated mice compared to control group (p<0.0001). **(C)** Levels of NE were measured by ELISA from the EV fraction isolated form BALF of the LPS treated and control mice and EV-depleted BALF of all mice. ELISA showed that LPS-induced lung inflammation increases the levels of both EV-associated NE and free NE in the BALF of LPS treated as compared to control mice (p<0.05). **(D)** LPS-induced lung inflammation resulted in increased EV-associated NE in the plasma of LPS treated mice (p<0.05) while no significant differences was observed in the levels of free NE in EV-depleted serum between two groups (p=0.14).

### *In vivo* Biodistribution of DIR Labeled EV

We hypothesized NE^+^EV from inflamed lungs enter the bloodstream and reach the liver. To test this, NE^+^EV were isolated from mouse bone marrow-derived neutrophils and labeled with DIR, a lipophilic fluorescent dye. We delivered 1×10^9 DIR-labeled NE^+^EV intratracheally to lungs of wildtype C57BL/6 mice with LPS-induced lung inflammation. The same volume of supernatant of the EV fluorescent labeling process was used as control. Imaging was performed using the IVIS® system at 6 and 24 hours (n=3/group). Figure 3A shows typical IVIS images of the *in vivo* biodistribution of dye and NE^+^EV. High accumulation of labeled EVs was observed in the liver area for both samples, consistent with previous data (20). We also delivered 1×10^9 DIR-labeled NE^+^EV, NE^-^EV and precipitated dye intravenously through tail vein injection to wildtype C57BL/6 mice and performed imaging on the harvested organs after 24 hours (n=3/group). Figure 3B shows IVIS images of *ex vivo* biodistribution of dye, NE^-^EV and NE^+^EV. High accumulation of injected EVs was observed in the liver and spleen for both EV-injected groups. (20). *Ex vivo* imaging demonstrated the liver uptakes the majority of intravenously injected EVs. Interestingly, we observed a higher DIR signal from the liver of mice injected with NE^-^EV compared to those injected with NE^+^EV. However, quantification of the radiant efficiency revealed no significant difference between the groups (Figure 3C). These results confirm NE^+^EV from inflamed lungs can reach and accumulate in the liver, supporting the hypothesis that circulating NE^+^EV contribute to liver pathology.

**Figure 3.**
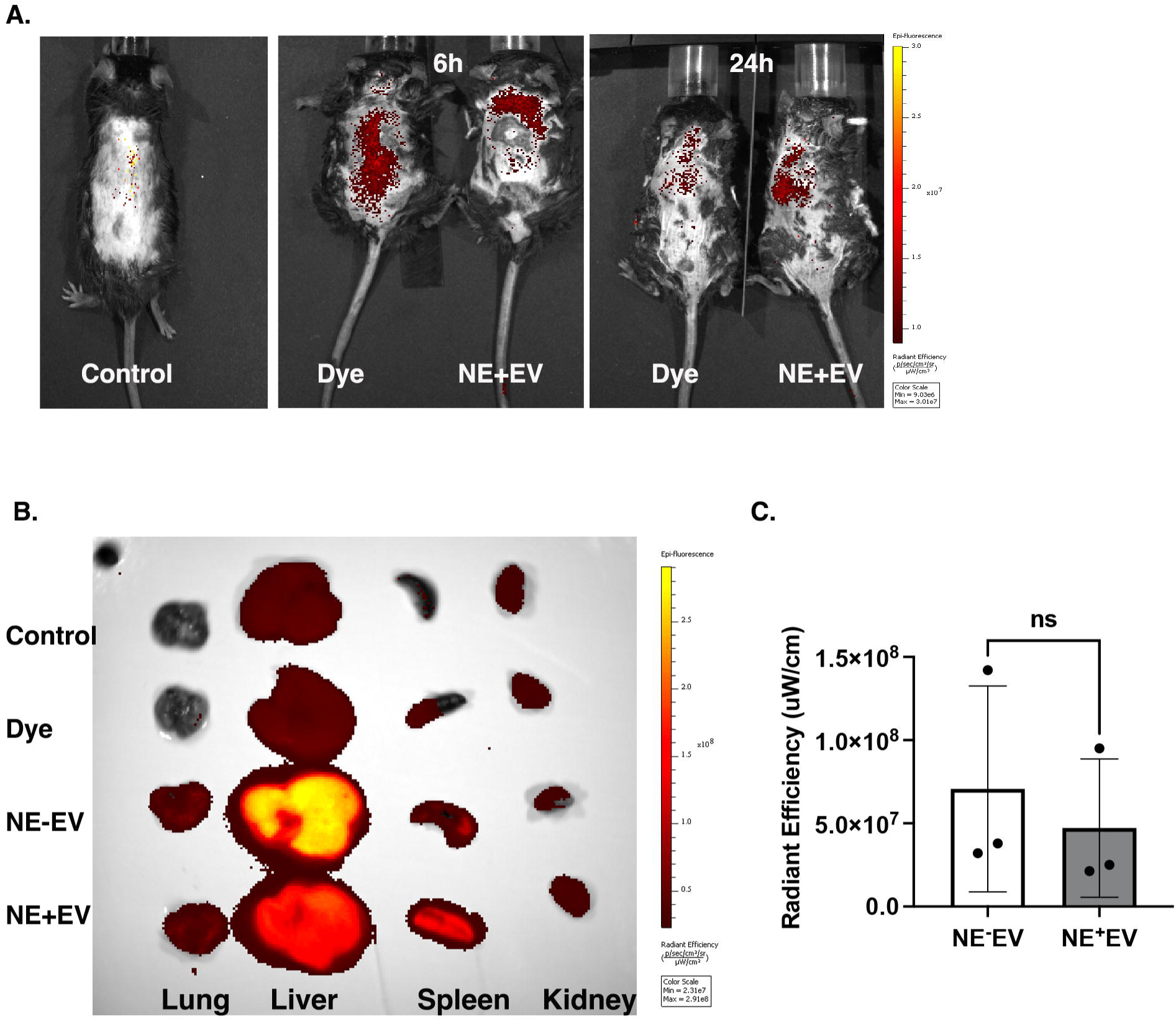
DIR-labeled NE^-^EV and NE^+^EV from blood circulation are up taken by the liver *in vivo*. **(A)** NE^+^EV isolated from neutrophils and labeled with DIR fluorescent dye. 1×10^9^ DIR-labeled NE^+^EV were intratracheally delivered to the lung of wildtype mice with LPS-induced lung inflammation. The same volume of precipitated dye was used as control. Imaging was performed using the IVIS® system and showed high accumulation of fluorescent signal in the liver area of the mice both at 6 and 24h. **(B)** 1×10^9^ DIR-labeled NE^+^EV, NE^-^EV and precipitated dye were intravenously delivered to wildtype C57BL/6 mice and imaging was performed on the harvested organs after 24h. Results of the *ex vivo* biodistribution of dye, NE^-^EV and NE^+^EV showed high accumulation of fluorescent signal in the liver and spleen. **(C)** Quantification of the radiant efficiency revealed no significant difference between DIR signal from the liver of the mice injected with NE^-^EV as compared with NE^+^EV (p>0.05).

### NE^+^EV Induce Activation of LX2 Cells

Light microscopy was employed to assess the impact of NE^+^EV on the morphology of LX2 cells. Following 24 hours of NE^+^EV treatment, over 50% of LX2 cells exhibited notable morphological changes. These changes included gathering, clumping and stretching, indicative of the activation state of the cells (Figure 4A). This structural reconfiguration suggests NE^+^EV treatment prompts a transition in LX2 cells towards an activated phenotype. To further evaluate fibrotic effects of NE^+^EV on LX2 cell activation, we analyzed RNA expression levels of α-SMA and Col1A1, key fibrogenesis markers, using quantitative PCR (qPCR). Compared to the control and NE^-^EV treated cells, NE^+^EV treatment significantly increased expression levels of α-SMA (Figure 4B) and Col1A1 (Figure 4C). The elevated expression of these fibrogenic markers indicates EV-associated NE effectively induces activation of LX2 cells. These findings collectively demonstrate NE^+^EV treatment not only alters the morphology of LX2 cells, but also enhances expression of genes associated with fibrogenesis, promoting activation of hepatic stellate cells.

**Figure 4.**
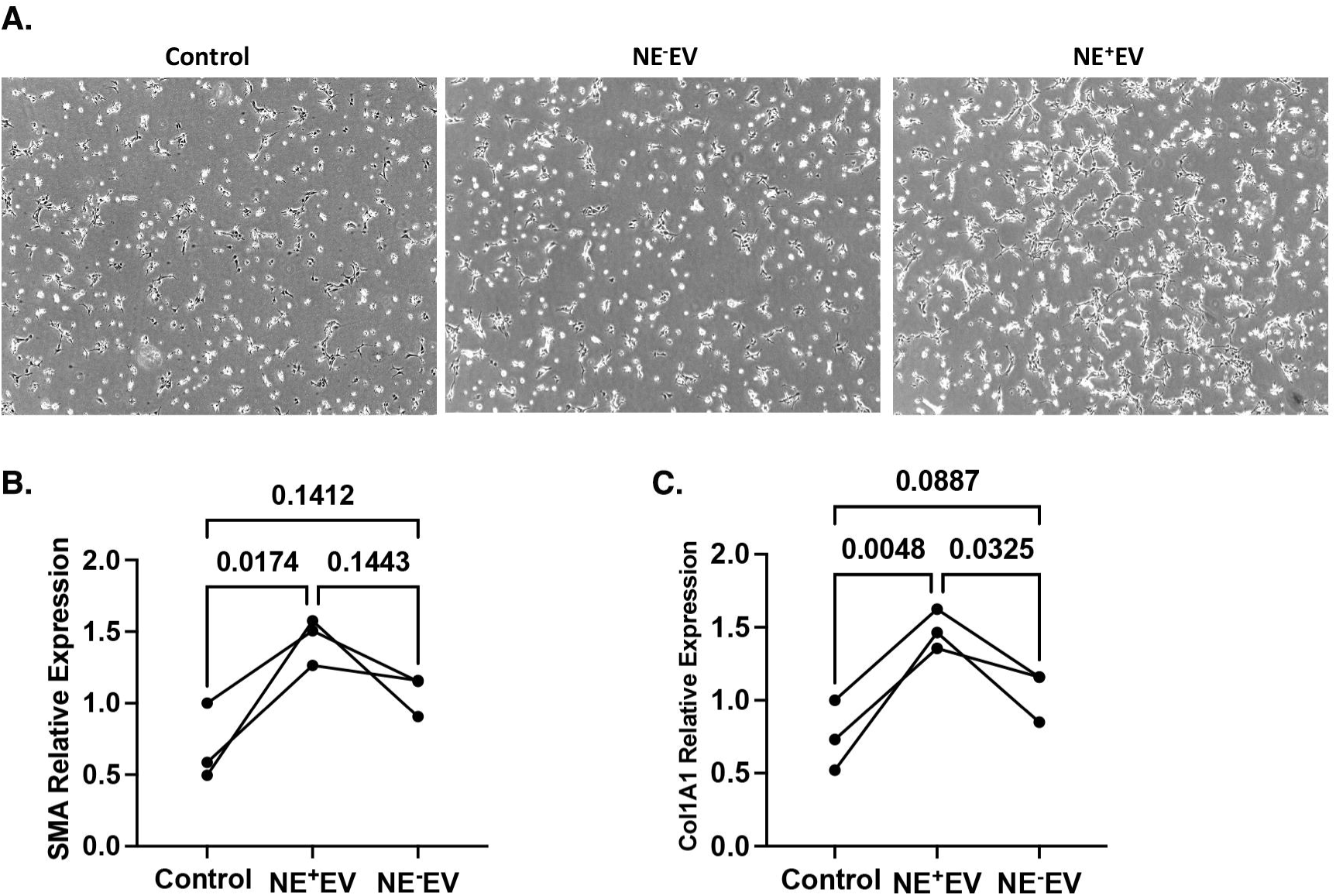
NE^+^EV treatment mediates changes in morphology and activation state of LX2 cells. **(A)** Following 24 hours of NE+EV treatment of LX2 cells, light microscopy revealed over 50% of LX2 cells exhibited significant morphological changes toward activation state. The morphological changes include gathering, clumping and stretching of the cells. **(B)** Quantitative PCR (qPCR) analysis was conducted to measure RNA expression of α-smooth muscle actin (α-SMA) in LX2 cells following treatment. Data from three independent experiments showed treatment with NE^+^EV significantly increased α-SMA gene expression compared to control groups and cells treated with NE^-^EV (p<0.05). **(C)** Similar qPCR analysis was performed to assess RNA expression levels of collagen type I alpha 1 chain (Col1A1) in LX2 cells. The results demonstrated a significant increase in Col1A1 gene expression in NE^+^EV treated cells compared to both control and NE^-^EV treated cells (p<0.05).

### NE^+^EV Increases Migration and Contraction Capability of LX2 Cells

To further validate the hypothesis that NE^+^EV promote activation and migration of LX2 cells, a wound healing assay was conducted over 30 hours. The results demonstrated NE^+^EV treatment significantly enhanced migration of LX2 cells into the wound area as compared to the control and NE^-^EV treated cells which showed a lesser extent of migration (Figure 5A). The difference in migration between NE^+^EV treated cells and control cells was statistically significant, highlighting the potent effect of NE^+^EV on migratory capabilities of LX2 cells (Figure 5B). These findings suggest NE^+^EV effectively promote the migratory ability of HSC *in vitro*. We further investigated whether NE^+^EV influence HSC contraction by employing a gel contraction assay using LX2 cells. The assay results indicated NE^+^EV treatment markedly induced gel contraction by HSC, with a significant difference observed as early as 24 hours post-treatment (Figure 5C). In contrast, control cells and NE^-^EV-treated cells exhibited minimal contraction, which was not significantly different from each other (Figure 5D). This enhanced contraction suggests NE^+^EV not only activate but also enhance the contractile properties of HSC. Collectively, these data confirm NE^+^EV treatment positively contributes to both migration and contraction of HSC in vitro, supporting the role of NE^+^EV in promoting key fibrogenic behaviors of these cells (Figure 5E, BioRender.com).

**Figure 5.**
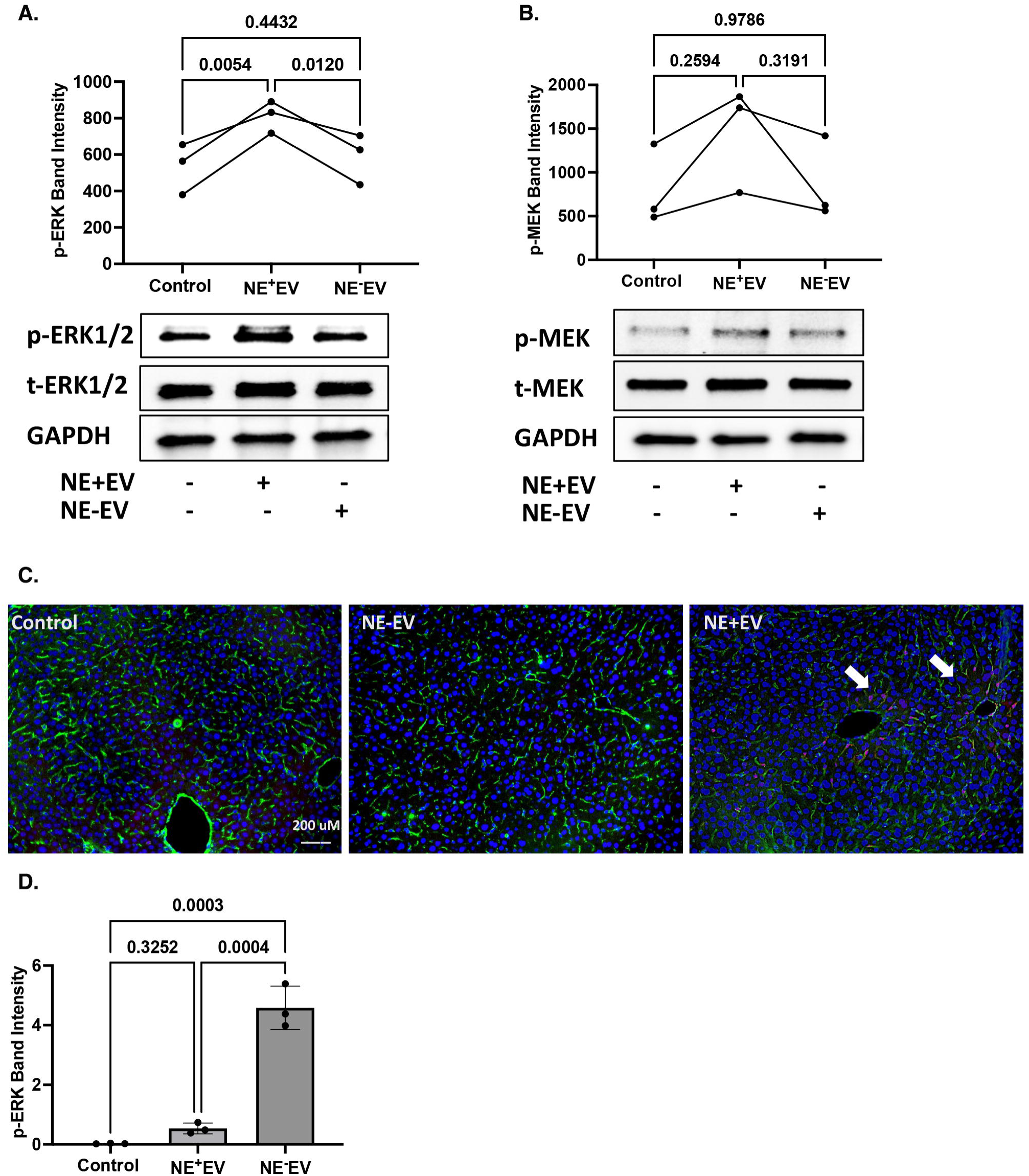
NE^+^EV treatment results in increased migration and contractile ability of LX2 cells. **(A)** Wound healing assay was performed for 30 hours using LX2 cells treated with NE^+^EV and NE^-^EV. LX2 cells were plated at 500,000 cells/well and the cell-coated surface was scraped with a 200μL pipette tip in a single stripe. The cultures were allowed to heal for 30h at 37°C. Migration of cells into wounded areas was photographed. Results show that NE^+^EV treatment aggravated migration of LX2 cells into the wounded area, while incubation of LX2 cells with NE^-^EV had lesser effects on migration as compared to control cells. **(B)** Graph showing the number of the cells migrated into the wounded area in each cell culture condition (n=3 independent experiments). **(C)** Gel contraction assay was performed using LX2 cells cultured on a Matrigel and treated with NE^+^EV and NE^-^EV for 24 hours. Results showed that compared to control LX2 cells and cells treated with NE^-^EV, NE^+^EV markedly induced LX2 gel contraction at about 24h treatment. **(D)** Graph shows percentage of the gel contraction calculated from 3 independent experiments (p<0.05). **(E)** Schematic picture showing NE^+^EV promoted HSC migration and contraction *in vitro* (BioRender.com).

### EV-associated NE Activates ERK1/2 Signaling Pathway in LX2 Cells

We investigated whether EV-associated NE could activate the ERK1/2 signaling pathway in LX2 cells. Our results showed NE^+^EV treatment significantly enhanced phosphorylation levels of ERK1/2 and MEK proteins in LX2 cells after just 6 minutes of incubation compared to the control group and cells treated with NE^-^EV (Figure 6A, 6B). To confirm these findings in vivo, we administered NE^+^EV intravenously to wild-type mice and observed phosphorylation of ERK1/2 in HSC 6 hours post-injection. Immunofluorescence analysis showed a strong red signal indicating phosphorylated ERK1/2 in HSC, with vimentin (a stellate cell marker) shown in green and nuclei in blue (Figure 6C, 6D). To confirm the pathologic role of NE on the surface of EV derived from activated neutrophils, we pre-incubated LX2 cells with Sivelestat, a neutrophil elastase inhibitor, before treating them with NE^+^EV. Western blot analysis revealed Sivelestat treatment significantly reduced phosphorylation of ERK1/2 and MEK proteins induced by NE^+^EV (Figure 7A). Band intensity quantification of phosphorylated ERK (p-ERK) and phosphorylated MEK (p-MEK) supported these results (Figure 7B, 7C). These results align with literature indicating NE induces activation of the MAPK signaling pathway (21) and suggests that EV-associated NE positively affects key signaling pathways in HSC. This evidence highlights the significant impact of NE^+^EV on the activation and migration of HSC, thereby promoting liver fibrogenesis.

**Figure 6.**
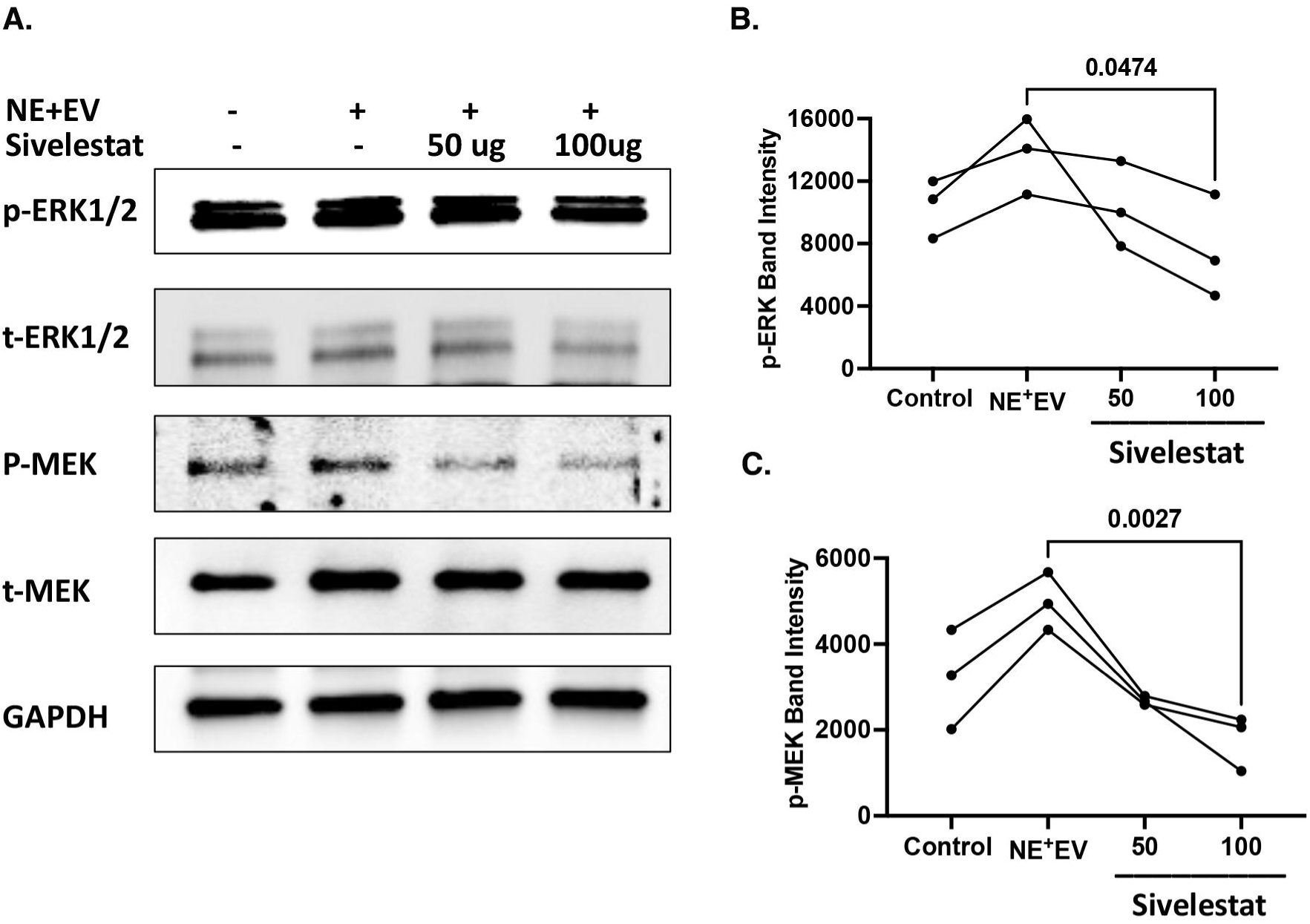
NE^+^EV mediates activation of LX2 cells through ERK1/2 signaling pathway. **(A)** Phosphorylation of ERK1/2 was assessed 6 minutes after treatment of LX2 cells with and without NE^-^EV and NE^+^EV using western blot analysis. Compared to non-treated control group and LX2 cells treated with NE^-^EV, NE^+^EV treatment markedly enhanced levels of phospho-ERK1/2 in LX2 cells. Quantification of band intensity from 3 independent experiments showed significant increases in the phospho-ERK1/2 band from NE^+^EV treated LX2 cells. GAPDH was used as loading control. **(B)** Phosphorylation of MEK, downstream of ERK1/2, was assessed in LX2 cells after 6 minutes of incubation with NE^-^EV and NE^+^EV and the result revealed higher phospho-MEK in cells treated with NE^+^EV as compared with NE^-^EV and negative control. **(C)** We intravenously administered NE^-^EV or NE^+^EV to wildtype mice and assessed the phosphorylation of ERK1/2 in hepatic stellate cells 6 hours post injection in the liver of the mice by immunofluorescent staining. We observed higher levels of red fluorescent signal (594nm) indicating phospho-ERK1/2 co stained with vimentin (488nm) as indicated by green signal in the liver of mice injected with NE^+^EV (n=3). **(D)** Graph shows red signal intensity in the liver of control, NE^-^EV and NE^+^EV injected mice, indicating significantly increased red signal in the images from NE^+^EV injected mice (p<0.05).

**Figure 7.**
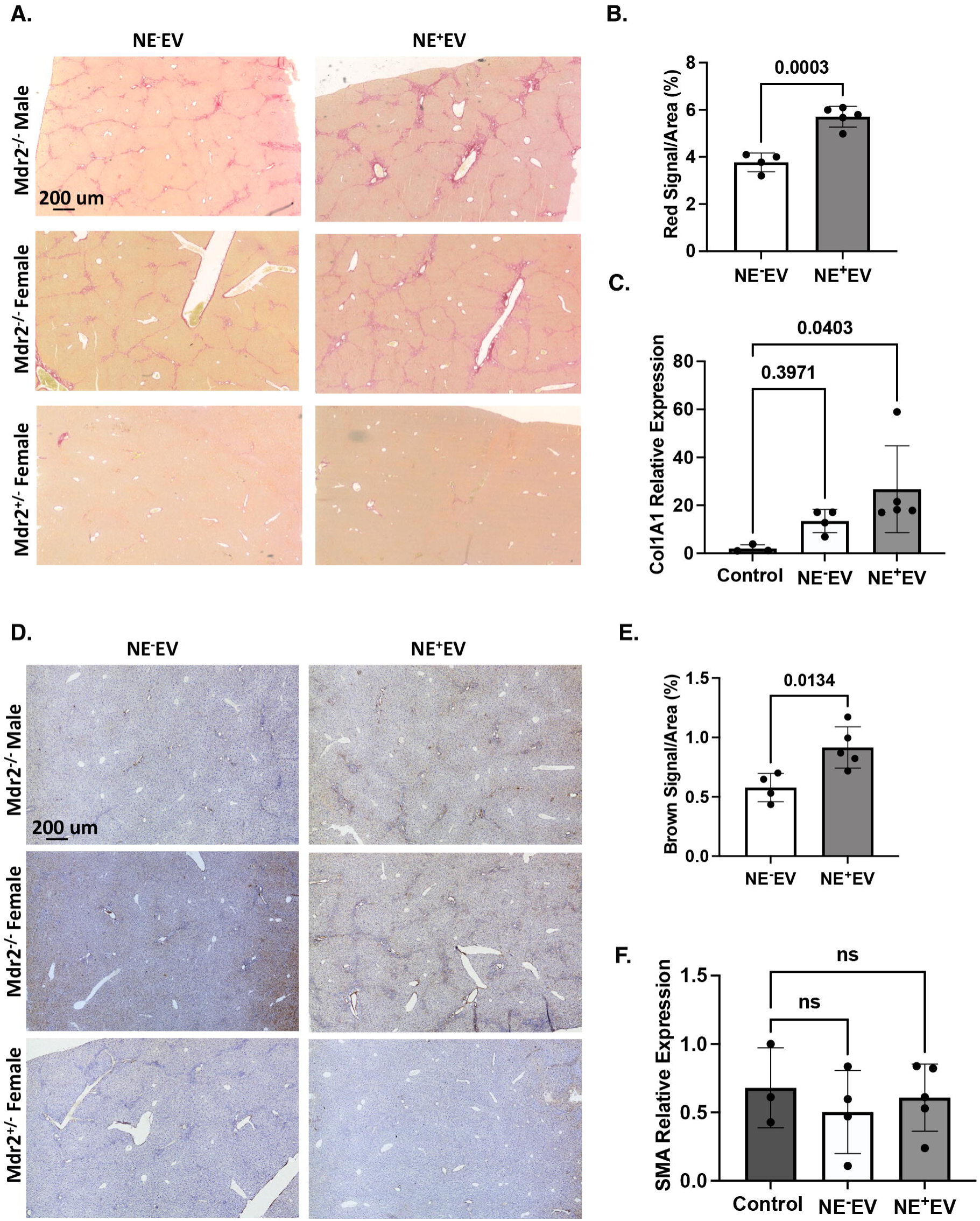
NE^+^EV-mediated activation of ERK1/2 pathway in LX2 cells depends on NE. **(A)** Western blot analysis showed incubation of LX2 cells with Sivelestat, a neutrophil elastase inhibitor, before treatment with NE^+^EV results in the reduction of the phosphorylated ERK1/2 and MEK proteins in LX2 cells as mediated by NE^+^EV. **(B)** The band intensity quantification of p-ERK and **(C)** p-MEK indicated significant decrease in the p-ERK and p-MEK signal when NE on the surface of EV was inhibited using Sivelestat treatment in LX2 cells, indicating the role of EV-associated NE in activating LX2 cells via activation of ERK1/2 pathway.

EV-associated NE Aggravates Liver Fibrosis in Mdr2^-/-^ Mouse Model of Liver Fibrosis The Mdr2^-/-^ mouse model of sclerosing cholangitis, characterized by impaired biliary phospholipid secretion and bile acid accumulation, develops spontaneous liver fibrosis by 6 weeks of age (22). We utilized this liver injury model to investigate the pro-fibrotic effects of EV-associated NE *in vivo*. We employed picrosirius red, α-SMA and Masson’s Trichrome staining to evaluate the fibrogenic role of EV-associated NE. In supplementary Figure 1, liver slices from healthy controls (Mdr2^+/-^) showed normal structural integrity. However, NE^+^EV administration in the experimental group (Mdr2^-/-)^ resulted in significant ECM deposition and loss of structural integrity. The NE-EV treated group showed less fibrosis and ECM deposition compared to the NE^+^EV group. Masson’s Trichrome staining (supplementary Figure 1) revealed blue-stained collagen in fibrotic tissue. Similarly, picrosirius red staining demonstrated significantly increased collagen I (Figure 8A, 8B) and α-SMA was also increased (Figure 8D, 8E) in the NE^+^EV treated group, evident as red and dark brown staining, respectively. NE^+^EV treated Mdr2^-/-^ mice showed a larger area and stronger staining with more positive cells compared to controls (p < 0.0003 and 0.0134). The healthy normal group (Mdr2^+/-^) showed minimal staining (p > 0.05). qPCR results supported these findings, indicating increased Col1A1 gene expression in Mdr2^-/-^ mice injected with NE^+^EV compared to those injected with NE^-^EV (Figure 8C). However, no significant differences in α-SMA gene expression were detected between the groups (Figure 8F). These results collectively demonstrate EV-associated NE exacerbates liver fibrosis in the Mdr2^-/-^ mouse model, significantly promoting ECM deposition and fibrogenic marker expression *in vivo*.

**Figure 8.**
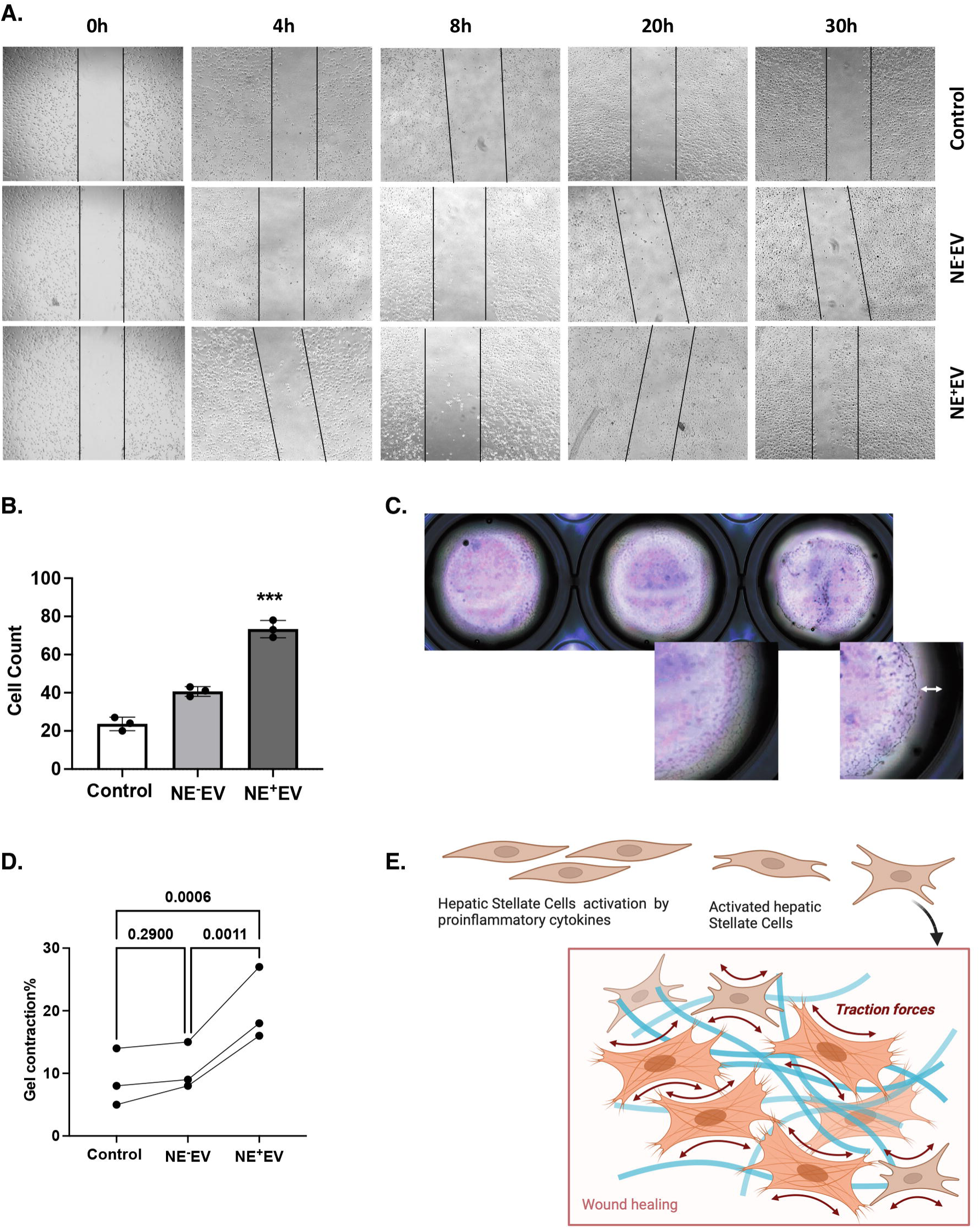
NE^+^EV exacerbate liver fibrosis in Mdr2^-/-^ mouse model of liver fibrosis. **(A)** Mdr2^-/-^ mouse model of sclerosing cholangitis was used to investigate the pro-fibrotic effect of EV-associated NE *in vivo*. Mdr2^-/-^ mice were injected intravenously with and without NE^-^EV and NE^+^EV for 3 weeks (3 times/week). After 3 weeks the liver tissues were harvested from the mice and analyzed. Picrosirius Red was used to evaluate the fibrogenesis role of EV-associated NE. **(B)** The results from Picrosirius Red staining from the liver tissues demonstrated that compared to NE^-^EV treated mice, the experimental mice injected with NE^+^EV have significantly increased signal from collagen I indicated as red signal. The staining in the liver of the Mdr2^-/-^ mice treated with NE^+^EV showed a larger area and stronger staining and more positive cells were observed compared with the control groups (*p* < 0.0005). The staining of collagen I in the normal group was not remarkable (*p* > 0.05). **(C)** The qPCR results from the same liver tissues indicated increased expression of the Col1A1 gene in Mdr2^-/-^ mice injected with NE^+^EV as compared with Mdr2^-/-^ mice injected with NE^-^EV. **(D)** α-SMA staining was used to evaluate the fibrosis progression in the experimental mice. **(E)** The results from α-SMA staining also demonstrated that compared to the NE^-^EV treated mice, the mice injected with NE^+^EV have significantly increased signal from α-SMA indicated as brown signal (*p* < 0.05). **(F)** Despite the fact that the liver of the mice injected with NE^+^EV had more α-SMA signal, we were not able to detect significant differences in the gene expression of α-SMA gene between the groups by qPCR analysis (p>0.05).

## Discussion

Here, we demonstrate lung inflammation increases plasma levels of EV-associated NE in mice, which reach the liver and promote HSC activation *in vivo*. Furthermore, incubation of LX2 cells with NE^+^EV mimics induced ECM synthesis by HSC and upregulates α-SMA and Col1A1 gene expression through the ERK/MAPK signaling pathway. In addition, we found intravenous administration of EV-associated NE aggravates liver fibrosis in the Mdr2^-/-^ mouse model of liver fibrosis, suggesting inflammation-derived EV-associated NE is, at least partially, involved in progression of liver fibrosis in pre-existing liver disease conditions. This study reveals an unappreciated aspect of organ crosstalk and interplay between systemic inflammation and liver disease with potential implications for future translational research.

Lungs are the major tissue reservoir for neutrophils and neutrophilic inflammation is a primary lung response to inflammatory challenges (23). During lung inflammation, NE^+^EV released by activated neutrophils have been observed to disrupt lung epithelial barriers and are found in the bloodstream (15, 24, 25). Previous studies show higher levels of NE^+^EV in mouse BALF and plasma during lung inflammation (15, 19). Our data also demonstrated LPS-induced lung inflammation increases the levels of both free and EV-associated NE in the BALF of mice with lung inflammation. However no significant effect was observed on the levels of free NE in the plasma of the mice in response to lung inflammation while higher levels of EV-associated NE was found in the plasma of LPS-treated mice. The activity of free NE has been shown to be stably inhibited by α1-antitrypsin that covalently binds to NE. However, EV-associated NE has intact enzymatic activity and is resistant to α1-antitrypsin (14, 15). Consistent with our previous work (26), these findings suggest increasing levels of plasma circulating proteolytic EV-associated NE during lung inflammation could ultimately affect the fate of other tissues and contribute to subsequent organ dysfunction.

The ability to track and measure EV biodistribution *in vivo* is essential for studying EV. Previous studies demonstrated tissue distribution of injected EV in mice, indicating the liver shows a strong propensity for up taking EV within a span of 2 to 24h (27). We employed DIR-fluorescent labeling to track neutrophil-derived EV, revealing DIR-labeled EV administered to the lungs primarily accumulate in the liver with a lesser amount in the spleen. Our *ex vivo* experiments further demonstrated both DIR-labeled NE^-^EV and NE^+^EV are predominantly taken up by liver tissue from the bloodstream. Notably, liver tissues of mice injected with NE^-^EV exhibited a higher DIR signal compared to those injected with NE^+^EV. This suggests NE^+^EV may experience greater fusion and uptake by recipient cells, potentially leading to diffusion of the fluorescent signal and a lower apparent concentration in liver tissue. However, the differential uptake rates and mechanisms of NE^-^EV versus NE^+^EV by liver cells warrant further investigation. These results support our hypothesis that EV derived from lung activated neutrophils can reach the liver via the bloodstream. This underscores the significance of examining how circulating inflammatory EV influence liver pathology, particularly in the context of systemic inflammatory diseases.

Liver fibrosis, resulting from progressive production and accumulation of ECM, is primarily caused by the activation of HSC, the most critical step in the development of liver fibrosis (28). HSC play a key role in initiation and progression of fibrogenesis in the liver by expressing and secreting collagen and α-SMA and thereby propagating liver fibrosis (29). Most liver diseases leading to liver fibrogenesis are caused by transition of HSC from a quiescent to an activated state. Liver HSC have been shown to express receptors for several cytokines and other inflammatory mediators such as NE (3, 30). Protease-mediated activation of HSC represents a paradigm in the pathogenesis of liver fibrosis. However, accumulating studies have demonstrated that NE, a serine protease released by neutrophils, promotes activation of HSC and contributes to liver fibrogenesis (31, 32). Here, NE^+^EV treatment of LX2 cells led to changes in cell morphology, an increase in gene expression levels of α-SMA and COL1A1, and increased migration and contractile ability. Nonetheless, NE^-^EV derived from activated neutrophils had lesser effects on the activation state of LX2 cells. While EV cargo is considered to be the same in NE^+^EV and NE^-^EV, these results indicate NE^+^EV exerts its promoting effect on HSC activation via NE on the surface of neutrophil-derived EV. This comprehensive analysis underscores the significant impact of NE^+^EV on HSC functionality, elucidating their role in liver fibrogenesis.

The MAPK signaling pathway is a key cascade integrating extracellular signals from cell surface receptors to regulate multiple cellular proteins involved in cellular activation (33). Previous studies show an aberrant MAPK signaling pathway contributes to development of organ fibrosis, such as in the liver. MAPK/ERK1/2 in particular plays an important role in cellular proliferation, adhesion, migration and survival to support liver fibrogenesis (34). NE activates the ERK1/2 pathway via activating transforming growth factor-α, epidermal growth factor receptor or protease activating receptors on the surface of different cell types (35). Similarly, our results demonstrated NE^+^EV stimulates phosphorylation of ERK1/2 and MEK in LX2 cells; this effect of NE^+^EV was inhibited with Sivelestat, a NE inhibitor. Our data shows in addition to the role of free NE released by liver infiltrated neutrophils in liver fibrogenesis, extrahepatic EV-associated NE also promotes HSC activation via positive regulation of the ERK/MAPK signaling pathway.

Normal liver is an immune-tolerant organ despite being exposed to various foreign molecules carried by the portal vein and systemic blood (36). However, several studies show injured liver fails to introduce a proper response to inflammatory stimuli or properly regulate immune mechanisms (37). In this regard, we used an Mdr2^-/-^ mouse model of liver fibrosis to test the profibrogenic effect of EV-associated NE on the injured liver *in vivo*. The Mdr2^-/-^ mouse model of sclerosing cholangitis develops spontaneous liver fibrosis at 6 weeks of age as a result of impaired biliary phospholipid secretion and bile acid accumulation (22, 38). Our experimental results confirmed IP injection of NE^+^EV exacerbates the process of liver fibrosis in the Mdr2^-/-^ mouse model of chronic liver injury as compared to NE^-^EV. Interestingly, NE^+^EV had no pro-fibrotic effect on the healthy liver of Mdr2^-/+^ mice. This suggests increased levels of circulating NE^+^EV might be associated with aggravation of liver injury and liver fibrosis in livers with pre-existing injury though not healthy liver.

According to previous studies, plasma levels of NE are associated with liver fibrosis in MASLD (39). Consistent with the literature, we demonstrated EV-associated NE exacerbates liver fibrosis in Mdr2^-/-^ mice and induces activation of HSCs *in vitro*. These findings suggest in addition to the impact of locally produced NE by liver-infiltrated neutrophils on the HSC activation state, excessive and/or inappropriately generated systemic NE can also promote liver damage and fibrogenesis, particularly when secreted NE associates with membranes such as EV which are resistant to inhibition by plasma protease inhibitors such as α1-antitrypsin (14, 15). Understanding the role of inflammatory plasma EV-associated NE in liver fibrosis is useful work. Eliminating NE^+^EV effects on HSC might bring promising treatment strategies for liver fibrosis in patients with systemic inflammation.

## Conclusion

Our findings indicate plasma EV-associated NE induces HSC activation and promotes liver fibrogenesis in the Mdr2^-/-^ mouse model of liver fibrosis. Despite experimental evidence discussed above, we acknowledge the fibrotic effect of EV-associated NE on other liver fibrosis models, such as those induced by α-1 antitrypsin deficiency, CCl4 and thioacetamide, remains to be further studied. Our research also elucidates the molecular mechanism of EV-associated NE-induced liver fibrosis involves phosphorylation of ERK1/2 and MEK, leading to activation and migration of HSC and subsequent liver fibrogenesis. The specific cell surface receptor responsible for NE-mediated HSC activation, along with its detailed mechanistic pathways, remains to be fully characterized and warrants further investigation. We demonstrated plasma circulation of NE^+^EV induced by neutrophilic inflammation and activation of neutrophils exacerbates liver fibrosis in pre-existing liver disease conditions. Understanding the roles of plasma circulating inflammatory EV in HSC proliferation, liver inflammation and their further functions during liver fibrogenesis in different types of liver diseases is crucial for future research. This could provide a therapeutic approach for treatment of liver fibrosis in inflammatory diseases.

## Data Availability Statement

The original contributions presented in the study are included in the article/supplementary material, further inquiries can be directed to the corresponding author.

## Ethics Statement

The studies were reviewed and approved by Institutional Animal Care and Use Committee (IACUC) Committee of the University of Florida (IACUC202300000056, and IACUC20230000502).

## Author Contributions

NK, HZ, BM and MH designed the study. RO performed the experiments. RO, ZG, IP, LK and BM participated in the collecting and analyzing of the data. NK and RO drafted and revised the manuscript. NK, HZ and MH edited the manuscript. MB provided funding. NK and MB supervised the study. All authors approved the final version of the manuscript for publication.

## Abbreviations

BAL: Bronchoalveolar lavage
BALF: Bronchoalveolar lavage fluid
DAB: Diaminobenzidine tetrahydrochloride
ECM: Extracellular matrix
EV: Extracellular vesicle
HSC: Hepatic stellate cells
NE: Neutrophil elastase
NE^+^EV: Extracellular vesicles positive for neutrophil elastase
NE^-^EV: Extracellular vesicles negative for neutrophil elastase
TEM: Transmission electron microscopy

**Figure Supplementary 1. NE^+^EV exacerbate liver fibrosis in Mdr2^-/-^ mouse model of liver fibrosis. (A)** Masson’s Trichrome staining from the liver of control group (Mdr2^+/-^) showed normal liver structural integrity. However, administration of NE^+^EV to the experimental group (Mdr2^-/-^) resulted in exacerbated fibrosis and the loss of structural integrity. In the NE^-^EV treated group, the slices showed less fibrosis and less ECM deposition compared with the experimental group. In Masson’s Trichrome images, collagen is shown as blue signal in fibrotic tissue. **(B)** The results from quantification of the blue signal of Masson Trichrome staining demonstrated compared to the NE^-^EV treated mice, the mice injected with NE^+^EV have significantly increased signal (*p* < 0.05).

