## Supplementary figures and images for "Extracellular Vesicle-Associated Neutrophil Elastase Activates Hepatic Stellate Cells and Promotes Liver Fibrogenesis via ERK1/2 Pathway"

### Supplementary figure 1

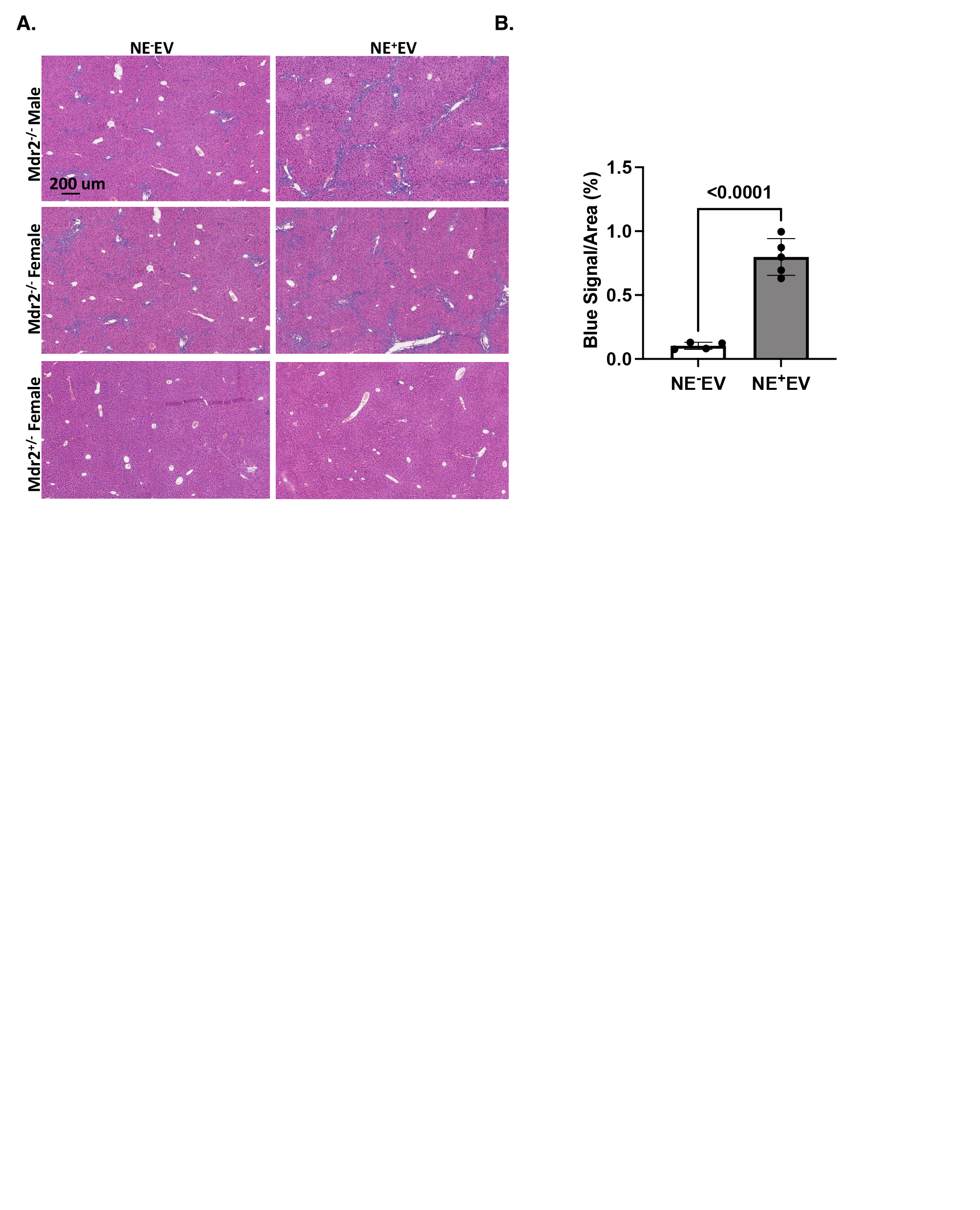
